# Sexual competition can select for male-only semelparity

**DOI:** 10.64898/2026.11.19.767966

**Authors:** Keaghan J Yaxley, Roberto Salguero-Gómez, Diana O Fisher

## Abstract

Semelparity – reproducing once in a lifetime – is an extreme solution to the trade-off between reproduction and future survival and is exceedingly rare in terrestrial vertebrates. Males of the Australian genera *Antechinus*, *Phascogale* and *Dasykaluta* are obligately semelparous while females are iteroparous, suggesting that the sexes face such different selective pressures that they have evolved different solutions to the same trade-off. Using invasion analyses of a novel two-sex population model with continuous reproductive investment, we show that the geometry of the reproduction–survival trade-off determines whether semelparity is evolutionarily accessible, but not necessarily whether it evolves. Age-structured survival and fecundity may maintain iteroparity under trade-off shapes for which classical theory predicts semelparity. Early onset of post-reproductive mortality can flip otherwise iteroparous trade-offs to semelparity, but only when survival declines quickly after sexual maturity. In contrast, when sexual competition gives high-investing males sufficiently disproportionate paternity, male-only semelparity can evolve under any trade-off geometry we examined, while females remain iteroparous. That male-only semelparity is rare while male competition is common suggests that lethal male investment evolves only under a particular combination of demographic, environmental and mating-system conditions.

## 1. INTRODUCTION

Semelparity – the life history strategy in which an organism reproduces only once during its lifetime – is an extreme solution to the trade-off that organisms make between current reproduction and future survival. The iteroparity-semelparity continuum describes the extent to which an organism disperses or concentrates reproductive effort across its lifetime (Fisher & Salguero-Gómez, 2026; Healy et al., 2019; Hughes, 2017). Obligately semelparous species are at the extreme of this continuum and are constrained to concentrate reproduction in one episode. For example, a hormonal trigger causes female octopus to inevitably cease feeding while brooding their only clutch, so they die of starvation but increase their reproductive success through maternal care (Wang & Ragsdale, 2018). Facultatively semelparous species are close to the semelparous extreme but are not constrained to concentrate reproduction in one episode. For example, female desert woodrats (*Neotoma lepida*) are typically iteroparous but a facultatively semelparous population occurs in one particularly hot location, where females that are large enough to raise large litters (increasing their fitness) die after reproduction because large size makes them vulnerable to extreme summer heat (Smith & Charnov, 2001). In a handful of taxa, only males are semelparous. For example, the male redback spider (*Latrodectus hasselti*) inserts himself into the female’s jaws, promoting sexual cannibalism in which he increases his paternity by extending copulation duration, increasing his lifetime fitness (Andrade, 1996; Forster, 1992).

Several theoretical explanations for the evolution of semelparous strategies have been posited. These can be divided into age-based demographic models, which focus on how mortality risk is distributed across an organism’s life (Charnov & Schaffer, 1973; Cole, 1954; Cushing, 2021; Murphy, 1968), and optimal allocation models, which focus on the dynamics of the trade-off between reproductive investment and future survival (Fisher & Salguero-Gómez, 2026; Gadgil & Bossert, 1970; Schaffer, 1974a, 1974b). Cole (1954) argued that most organisms should be semelparous because even a small increase in fecundity would increase fitness such that semelparity is favoured over iteroparity (Cole’s paradox). Charnov and Schaffer (1973) demonstrated that Cole’s paradox was a consequence of Cole’s model omitting age-specific differences in mortality rates, and showed semelparity was favoured when post-reproductive survival was low relative to juvenile survival. Further models have explored the impacts of environmental variability and density dependence (Benton & Grant, 1999; Orzack & Tuljapurkar, 1989) and interspecific competition (Cushing, 2021) on the evolution of semelparity.

Age structured models predict that semelparity will evolve when some biological or ecological driver lowers post-reproductive survival, such that an organism’s offspring have a better chance of surviving to reproduce than the adult has of reproducing again (Fisher & Salguero-Gómez, 2026). Optimal allocation models predict that the shape of the trade-off between age-specific reproductive effort and lifetime fitness will differ between semelparous and iteroparous taxa (Fisher & Salguero-Gómez, 2026). In semelparous organisms the fitness benefits of reproduction increase disproportionately at high levels of reproductive effort, or the costs of reproduction are initially high with low amounts of reproductive effort. In either case, the reproduction-survival trade-off is not concave – the relationship between the costs and benefits of increased reproductive effort are either linear or convex (Gadgil & Bossert, 1970; Schaffer, 1974a). Like the age-structured demographic models, the optimal allocation models predict that semelparous organisms should invest more in reproduction than their iteroparous relatives, but unlike the age-structured models this does not require an external driver of high post-reproduction mortality; that is instead a consequence of the reproduction-survival trade-off itself (Fisher & Salguero-Gómez, 2026).

Almost all animals have discrete sexes (∼94-95 per cent of described metazoans; Sasson & Ryan, 2017), and life history strategies often vary between the males and females (Ancona et al., 2020; Bonduriansky et al., 2008; Brooks & Garratt, 2017; Fairbairn, 1997). The most extreme divergence in male and female life histories among vertebrates is in the three genera of small Australian carnivorous marsupials (Family: Dasyuridae, *Antechinus, Phascogale* and *Dasykaluta*), in which the males are obligately semelparous and the females are iteroparous (Fisher et al., 2013). In A*ntechinus*, each year all females in a given location mate synchronously over a few days (Braithwaite & Lee, 1979; Collett et al., 2018). Both females and males mate with multiple partners. This compressed timing facilitates sexual competition and sexual conflict – males seek to monopolise paternity through mate-guarding mechanisms including prolonged copulation (∼12 hours), while females benefit from sperm competition because males with the most competitive sperm sire offspring with the highest survival, increasing lifetime fitness (Fisher, Double, Blomberg, et al., 2006). In dasyurids with obligate male semelparity, a surge in testosterone at sexual maturity triggers a cascade of escalating corticosteroid hormones, suppression of the immune system, internal haemorrhaging and infection. Within 2-3 weeks of the rut all males have died, while the females can reproduce again in the next season (Fisher & Blomberg, 2011). In addition to two clades of dasyurid marsupials, obligate male-only semelparity has evolved in seven known clades of spiders. Comparative analyses suggest that the driver in both mammals and arachnids is sexual selection involving sexual conflict, in which escalating male competition benefits female fitness (Fisher and Salguero-Gomez 2026).

To test if life history divergence fits the demographic or optimal allocation model, we need to find why drivers of life history evolution differ in males and females. In the first published adaptive explanation of semelparity in *Antechinus,* Braithwaite and Lee (1979) argued that semelparity in *Antechinus* results from their seasonally predictable environments, high post-reproductive mortality and the prolonged conception-to-weaning period of small marsupials. They argued that *Antechinus* are constrained to reproduce when young will be synchronously weaned in the month of annual peak prey abundance, which selects for semelparity as males are unlikely to survive 12 months to the next annual peak (the demographic model). A challenge for this argument is to explain why only males are semelparous whereas females can survive two seasons. Dickman (1994) argued that selection for intense sperm competition has driven the males of these species to trade-off post-reproductive survival for increased reproductive effort. In agreement, comparative analysis of insectivorous marsupial taxa (dasyurids and didelphids) identified sperm competition rather than post-reproductive mortality as the key predictor of male semelparity (Fisher et al., 2013). To determine whether sexual competition alone can decouple the life histories of males and females such that one is semelparous while the other is iteroparous, we need to understand the trade-offs in both sexes that underpin the optimal allocation model.

Here we explore whether sexual competition alone can act as a pathway to the evolution of semelparity in one sex of a species. We model the population dynamics of a small marsupial using a novel two-sex density-dependent matrix population model in which individuals in both sexes can invest differentially in reproduction, and where that investment trades off against future survival. Our model treats reproductive investment as a continuous trait, meaning that parity strategies under this model can fall along a continuum. We use this model to test three hypotheses. (H1) We first establish whether the predictions of optimal allocation hold using our model, i.e. that an iteroparous population is only evolutionarily viable when increased reproductive investment leads to diminishing fecundity returns or accelerating survival costs. We then test (H2) whether increasing the rate of post-reproductive male mortality for this stably iteroparous population will drive the males towards adopting a semelparous life history, as in the demographic model. Finally, we test (H3) whether a stable iteroparous population can be invaded by a semelparous male variant when higher-investing males sire a disproportionate share of offspring – i.e. can sexual competition without changes to extrinsic mortality, shift the male evolutionary optimum from iteroparity to semelparity while leaving the female strategy unchanged.

## 2. METHODS

### 2.1 – Population model and baseline vital rates

We modelled a hypothetical small marsupial population using a two-sex age-based structured population model that captures demographic dynamics and trade-offs, including aging (the transition of survivors through age classes) and reproduction (the production of recruits). The model tracked the male and female components of the population separately, each with its own set of vital rates and separate life history trade-offs. The male component was further subdivided into *k* male types, where each can be subject to a different set of vital rates. This allowed us to test whether mutant male life history strategies can invade an established resident male population, providing a framework for identifying evolutionarily stable strategies (ESS). This is necessary to evaluate the fitness of different male life history strategies, as male fitness is often zero-sum – they compete over the paternity of a restricted number of births limited by the number of females (Kokko & Brooks, 2003; Rankin et al., 2011). To ensure that our model reflects a biologically plausible demography for a small marsupial population, we used published demographic information for *Antechinus agilis* (Fisher, Double, & Moore, 2006) to inform its structure and parameterisation. The model and all analyses were implemented in the statistical programming language and environment R v.4.5.2 (R Core Team, 2025). The generative AI Claude (Anthropic; Opus and Fable models, accessed January–November 2026) was used to help write, revise and debug code for some analyses. The population model, its parameterisation, the analytical design and the interpretation of the results were all developed by the authors and the text was written by the authors. All AI tasks were directed, supervised, reviewed and verified by the corresponding author who takes full responsibility for their content.

The model is structured by discrete age classes *a*, spanning a maximum life span of 24 months consistent with the reported maximum lifespan of female *A. agilis,* such that *a* ∈ {0, 1, 2, …, *A*}, with *A* = 23 and step Δ*a* = 1. Throughout ℒ(*x*) = (1 + *e*^−*x*^)^−1^ denotes the standard logistic function. A semicolon inside a function’s arguments list separates its *arguments* – quantities to the left of the semicolon that vary as the model runs – from its *parameters* – quantities to the right of the semicolon that are fixed. A subscript ∗ ∈ {*m*, *f*} indicates that separate male and female versions of a quantity and *k* indexes male *m* investment types. The male and female components of the population at time *t* is given by the male and female age-distribution vectors

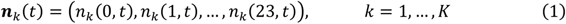

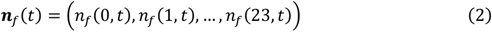

Where *n*_∗_(*a*, *t*) is the number of individuals of a given sex/type in age class *a* at time *t*. The total population size at time *t* is

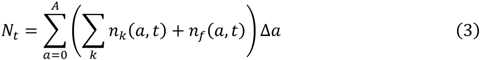

Table 1 gives a complete summary of all the model and analysis notation.

**Table 1.**
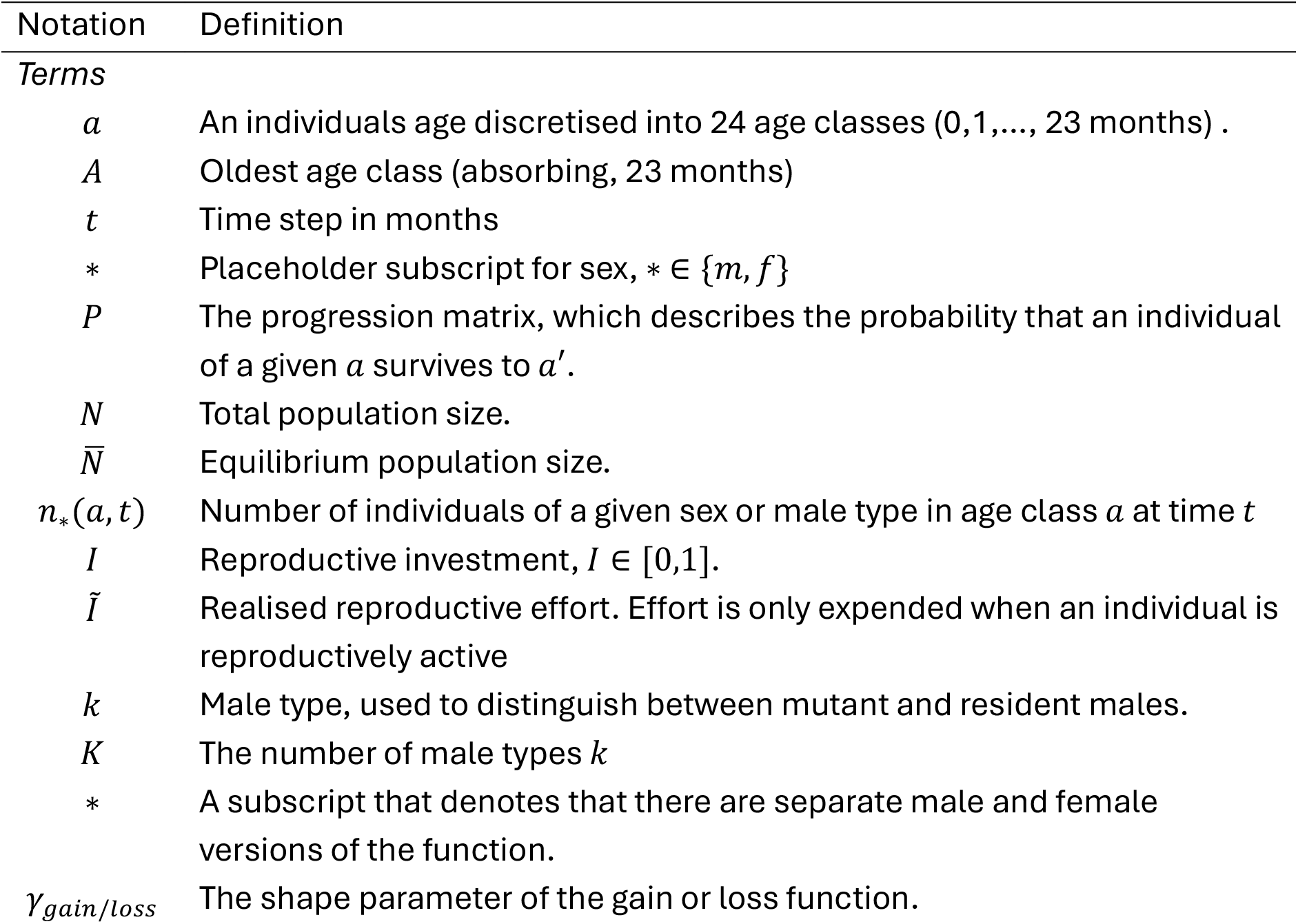
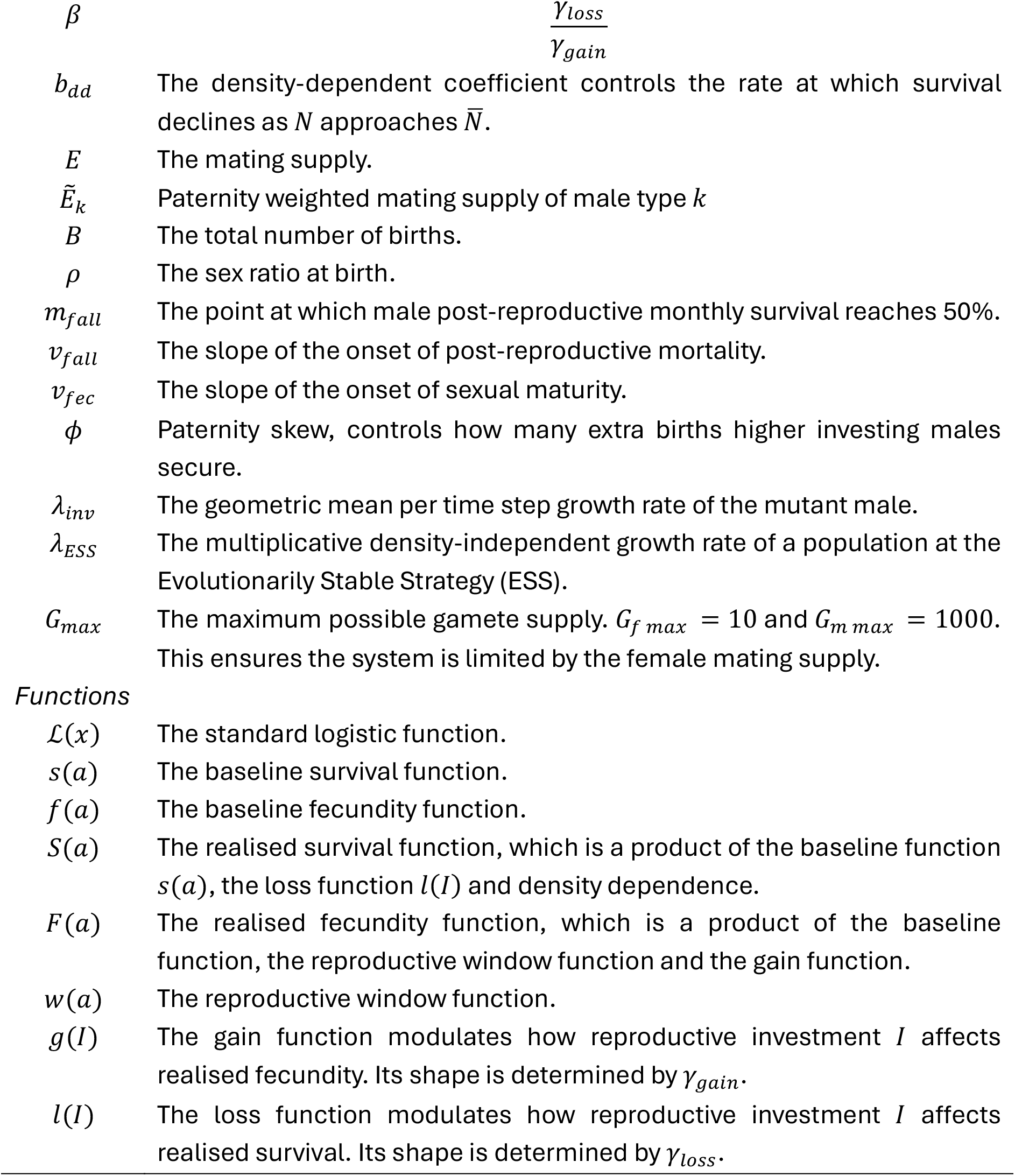
Notation used in the article.

| Notation | Definition |
|---|---|
| *Terms* |  |
| a | An individuals age discretised into 24 age classes (0,1,..., 23 months) . |
| A | Oldest age class (absorbing, 23 months) |
| t | Time step in months |
| ∗ | Placeholder subscript for sex, ∗ ∈ {m, f} |
| P | The progression matrix, which describes the probability that an individual of a given a survives to a′. |
| N | Total population size. |
| N̄ | Equilibrium population size. |
| n<sub>∗</sub>(a, t) | Number of individuals of a given sex or male type in age class a at time t |
| I | Reproductive investment, I ∈ [0,1]. |
| Ĩ | Realised reproductive effort. Effort is only expended when an individual is reproductively active |
| k | Male type, used to distinguish between mutant and resident males. |
| K | The number of male types k |
| ∗ | A subscript that denotes that there are separate male and female versions of the function. |
| γ<sub>gain/loss</sub> | The shape parameter of the gain or loss function. |
| β | $\frac{\gamma_{loss}}{\gamma_{gain}}$ |
| b<sub>dd</sub> | The density-dependent coefficient controls the rate at which survival declines as N approaches N̄. |
| E | The mating supply. |
| Ẽ<sub>k</sub> | Paternity weighted mating supply of male type k |
| B | The total number of births. |
| ρ | The sex ratio at birth. |
| m<sub>fall</sub> | The point at which male post-reproductive monthly survival reaches 50%. |
| v<sub>fall</sub> | The slope of the onset of post-reproductive mortality. |
| v<sub>fec</sub> | The slope of the onset of sexual maturity. |
| ϕ | Paternity skew, controls how many extra births higher investing males secure. |
| λ<sub>inv</sub> | The geometric mean per time step growth rate of the mutant male. |
| λ<sub>ESS</sub> | The multiplicative density-independent growth rate of a population at the Evolutionarily Stable Strategy (ESS). |
| G<sub>max</sub> | The maximum possible gamete supply. G<sub>f max</sub> = 10 and G<sub>m max</sub> = 1000. This ensures the system is limited by the female mating supply. |
| *Functions* |  |
| ℒ(x) | The standard logistic function. |
| s(a) | The baseline survival function. |
| f(a) | The baseline fecundity function. |
| S(a) | The realised survival function, which is a product of the baseline function s(a), the loss function l(I) and density dependence. |
| F(a) | The realised fecundity function, which is a product of the baseline function, the reproductive window function and the gain function. |
| w(a) | The reproductive window function. |
| g(I) | The gain function modulates how reproductive investment I affects realised fecundity. Its shape is determined by γ<sub>gain</sub>. |
| l(I) | The loss function modulates how reproductive investment I affects realised survival. Its shape is determined by γ<sub>loss</sub>. |

The different vital rates of the model are described by functions. We model age-specific baseline survival rates using a rise-fall logistic function and parametrised it with published survivorship data for female *A. agilis* (Fig 1; Supporting Information S1; Supporting Information S2; Equation S1).

**Figure 1.**
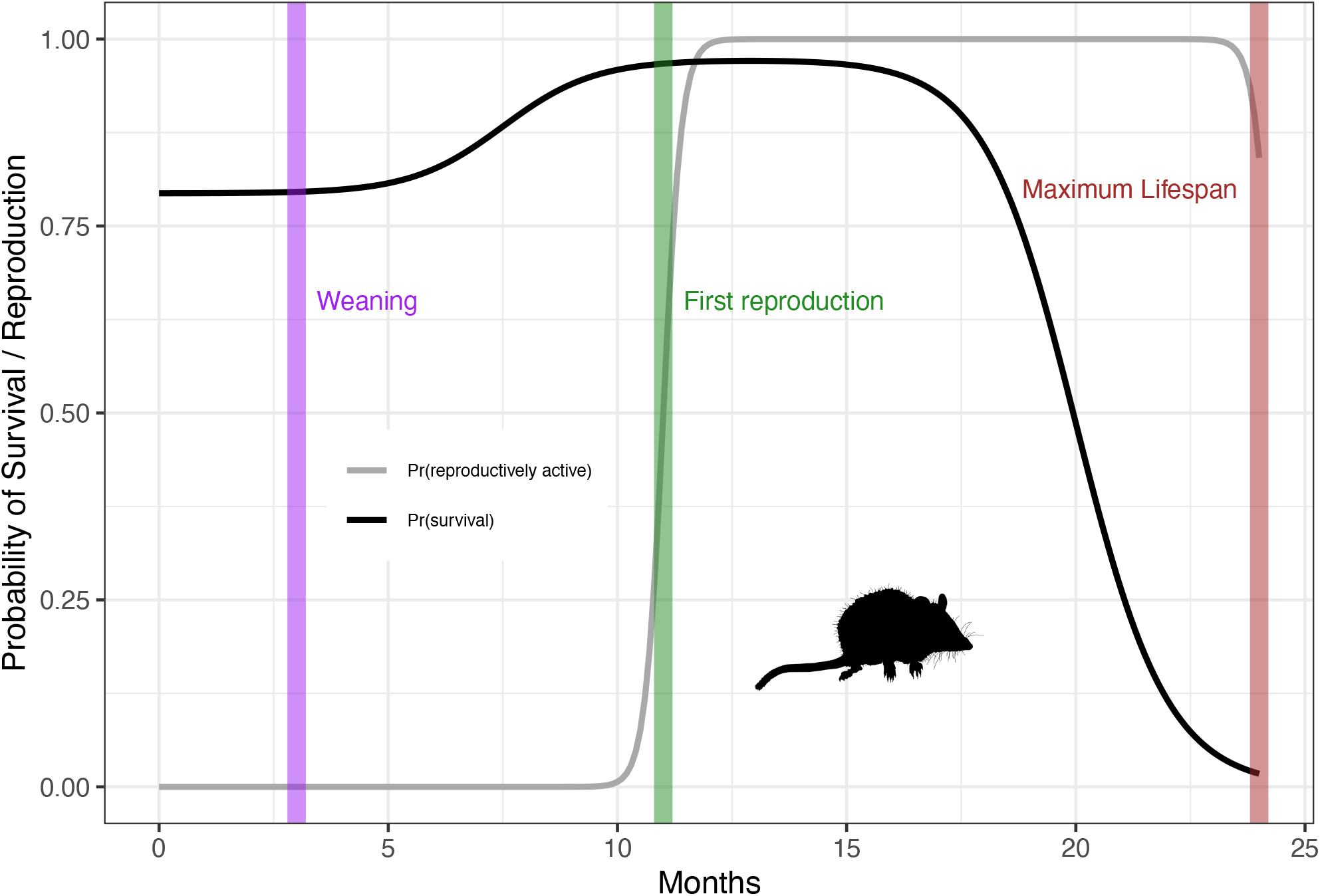
Survival (black) and reproductive window (grey) functions based on reported demographic values for female *Antechinus agilis* (silhouette). Vertical bars indicate the species’ age of weaning (purple, 3 months), sexual maturity (green, 11 months) and the maximum lifespan (red, 24 months).

We modelled baseline reproductive outcomes with two functions. The first is the reproductive window function *w*(*a*) which describes how the probability that an individual is reproductively active changes with age (Fig 1; Supporting Information S2; Equation S2). The model runs on monthly age-classes, so individuals of both sexes reproduce and incur the survival cost of reproduction in every month from sexual maturity to death. We deliberately keep the reproductive window open over this period, rather than impose a brief synchronised breeding season as observed in *Antechinus*, because imposing a narrow window would risk making semelparity an inevitable outcome of our model. A broad window makes the test more conservative – semelparity must evolve even though individuals have the opportunity to reproduce multiple times across their lifetime.

The second is our fecundity function *f*(*a*) for which we use the constant *G_max_* meaning that reproductive output of individuals does not vary with age outside of its interaction with *w*(*a*) (Supporting Information S2; Equation S3). For the female component *G_fmax_* = 10 consistent with the average litter size reported for *A. agilis* (Ferreira et al., 2026) and for the male component we set *G_mmax_* = 1000. This ensures that, under most treatments, our model’s births will be female limited, and that male investment will translate to competition over those limited births. This is a biologically reasonable assumption for a small marsupial population.

### 2.2 – Reproductive investment and modelling the reproduction-survival trade-off

In our model individuals can “choose” how much effort they invest into reproduction as a strategy. An individual’s strategy is the scalar *I*_∗_ ∈ [0,1] and because effort is only expended when reproductively active the *realised*, *age-specific* effort is

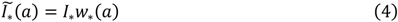

This allows us to model the trade-off between reproductive effort and future survival that defines the iteroparity-semelparity continuum (Healy et al., 2019; Hughes, 2017; Salguero-Gómez et al., 2016). The exact way *I*_∗_ affects the realised fecundity and survival of individuals is controlled via the gain function *g*(*I*_∗_) and the loss function *l*(*I*_∗_). The optimal allocation model predicts semelparity will be the evolutionarily stable life-history strategy when returns on reproductive effort are accelerating or the costs of reproductive investment to future survival are decelerating, or both (Gadgil & Bossert, 1970; Schaffer, 1974a, 1974b). Therefore, we define *g*(*I*) and *l*(*I*) as the monotonically increasing and decreasing power functions.

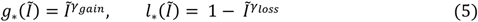

where *γ* controls the shape of the function. Because *l*_∗_(1) = 0 for any *γ_loss_*, maximum reproductive investment (*I* = 1) reduces survival to effectively zero once an individual is fully reproductively active (*w*(*a*) → 1). Therefore, *I* = 1 is treated as effectively lethal reproductive investment regardless of the trade-off geometry which only controls how rapidly survival declines as investment increases. We can describe the realised fecundity as baseline fecundity modulated by the gain function (Supporting Information S2; Equation S4) and realised survival.

Realised survival is described as baseline survival modulated by the loss function and negative-exponential density dependence (Supporting Information S2; Equation S5). A negative-exponential density dependence term ensures that as the total population size *N* increases, survival declines smoothly, and that eventually the population reaches an equilibrium size of 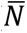. Density dependence ensures the *k* type males compete over a limited number of possible births. This is an important criterion for measuring male fitness – males participate in a zero-sum game over paternity (Kokko, 2021; Kokko & Brooks, 2003; Rankin et al., 2011). Survivors advance exactly one age class per time step and the final age class *A* is absorbing (Supporting Information S2; Equation S6).

### 2.3 – Two-Sex Mating Function

To determine the total number of births we first calculate the male mating supply *E_m_* and the female mating supply *E_f_*, which is the realised fecundity summed over age classes (Supporting Information S3; Equation S7). We then use the harmonic mean to determine the number of potential births *B* (Supporting Information S3; Equation S8). We use the harmonic mean because it is pulled towards the rarer of the two sexes’ mating supply, i.e. whenever either sex is very rare in the population *B* → 0. When mating supply is equal between the two sexes, then *B* = *E_m_* = *E_f_*. However, because females are the limiting factor in mammal reproduction, we scale the ratio of births by the total female mating supply and divide the paternity of these births between *k* type males such that each *B_k_* is proportional to each type’s raw mating supply, *E_k_* (Supporting Information S3; Equation S9). *B_k_* is then partitioned into male and female recruits based on a sex ratio *ρ* = 0.5 (*A. agilis* did not have a biased sex ratio at birth (Fisher, Double, Blomberg, et al., 2006)).

Male recruits inherit the *I_k_* of their fathers as if it were a patrilineal/Y-linked trait. This is a simplifying assumption that avoids the need to track autosomal genotype classes across both sexes and all age classes, but one that we do not expect to meaningfully affect the results of our invasion analyses. A paternal inheritance structure exposes every copy of the mutant allele to selection at every generation. This will accelerate the spread or loss of the mutant *I_k_* compared to an autosomal system but will not change the direction of selection on it (Supporting Information S4).

The complete model is

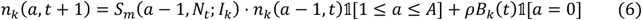

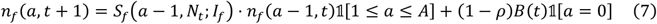

### 2.4 – Evolutionary invasion analysis

To test whether each of three pathways to semelparity can lead to male-only semelparity, we employ invasion analysis to identify the evolutionarily stable level of male investment. For a given set of gain and loss functions, we first project the resident population forward 500 time steps towards its density-dependent equilibrium (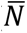 = 2000 for the ESS resident). We then introduce a rare mutant male with a different level of reproductive investment, at 0.0001% of male abundance, into the first age class (0 months). Mutant males differ from resident males in that mutants can vary in reproductive effort *I_k_*. The resident and mutant are projected 2,000 time steps forward. After discarding the first 500 timesteps as burnin, we record the mutant’s invasion fitness *λ_inv_* as the geometric mean per-time step growth rate of the mutant against the resident equilibrium, rescaling the mutant to its initial abundance after each step so that its age structure converges while its density remains negligible. If *λ_inv_* > 1 the mutant is increasing relative to the resident, and the resident strategy is therefore invasible. If *λ_inv_* < 1 the resident strategy is not invasible.

We use this invasion analysis in two ways. First, we create nine discrete gain-loss treatments based on each combination of an accelerating (*γ* = 2), linear (*γ* = 1), or decelerating (*γ* = 0.5) gain and loss functions, hereafter referred to as the gain-loss treatments (Figure 2). For these nine gain-loss treatments we compute *λ_inv_* for a given set of resident and mutant reproductive investment strategies, to test whether a mutant can invade a resident strategy. Female investment is solved jointly with male investment over the same grid but for all treatments the female gain function is decelerating (*γ* = 0.5) and the loss function accelerating (*γ* = 2). To check that the per-treatment calibration of the density-dependence coefficient *b_dd_* was not affecting the results of our invasion analysis, we ran a sensitivity analysis in which we halved and doubled *b_dd_* for the accelerating gain and linear loss and linear gain and accelerating loss treatments and recomputed invasion fitness (Supporting Information S5).

**Figure 2.**
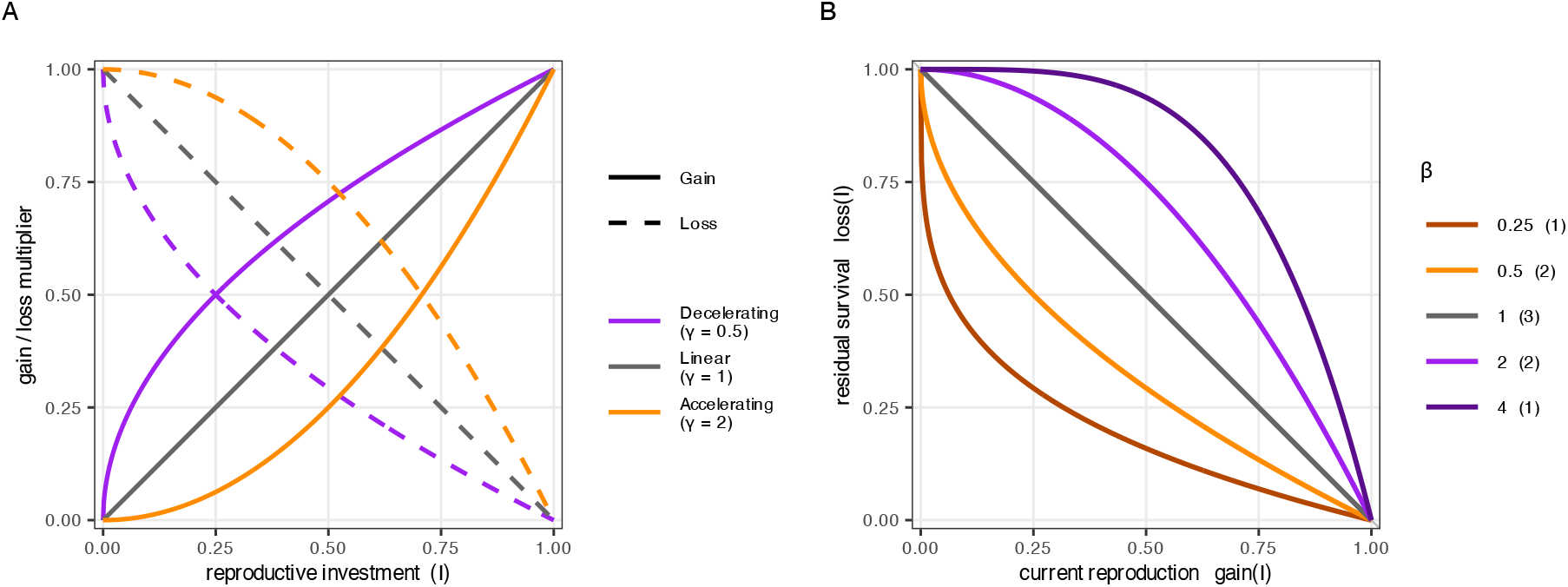
(**A**) Gain and loss functions used to modulate how reproductive investment (*I*_∗_) trades-off with fecundity and survival. These are symmetrical rising and falling power functions, with the shape determined by the parameter *γ*. (**B**) Curvature of the reproduction-survival trade-off for all possible combinations of the gain and loss functions in panel A. The colours of the lines indicate the value of *β* and the integer in brackets the number of gain-loss combinations that reduce to each value. Under the classical optimal allocation models, semelparity is expected to evolve when the trade-off curve is convex *β* < 1 or linear *β* = 1, and iteroparity will only be possible when the curve is concave *β* > 1.

Second, in our model the reproduction-survival trade-off is defined as two power functions (rising and falling), and so any combination of gain and loss functions can be collapsed into a single geometric quantity 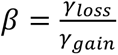 that governs the shape of the trade-off. The trade-off is convex when *β* < 1, linear when *β* = 1 and concave when *β* > 1. This is consistent with Gadgil and Bossert (1970) and Schaffer (1974b) models which predicts that when the reproduction-survival trade-off is convex or linear, semelparity will be the ESS, but when it is concave the ESS will be some intermediate amount of reproductive effort. We also employ our invasion analysis to identify the evolutionarily stable level of male investment across a continuous plane of the reproduction-survival trade-off, hereafter referred to as the full sweep analysis. Across all sweeps, female investment is modelled again with *γ*_*gain*_ = 0.5 and *γ_loss_* = 2 and *I_f_* = 0.30, as this is the ESS of the female trade-off given that combination of gain and loss parameters. In the full sweep analyses, resident and mutant investment take values *I* ∈ {0.1, 0.2, …, 1} so ESSs are located to ±0.05. To account for numerical error in the full-sweep analyses a mutant was counted as invading only if *λ_inv_* exceeded the invasion fitness of a mutant identical to the resident.

### 2.5 –H1: Testing the predictions of the optimal allocation model

To test the prediction of the **optimal allocation model**, that semelparity evolves when reproductive investment has accelerating fecundity returns or decelerating survival costs, we test whether the shape of the gain and loss function influences the invasibility of an iteroparous population by a semelparous male. For each of the nine gain-loss treatments we project our population forward across a grid of resident and mutant male investment levels *I* ∈ {0.05, 0.10, 0.15, …, 1}, to create a pairwise invasibility plot (PIP). We classified each singular strategy (or boundary strategy at *I_m_* = 1) as those that are evolutionarily stable (cannot be invaded by a mutant) and convergence stable (nearby residents can be invaded by mutants closer to it so selection moves the population towards it) (Geritz et al., 1998). We read both from the PIPs. We then additionally conduct a full sweep analysis by sweeping a 17 × 17 grid of *γ*_*gain*_ and *γ_loss_* spanning 0.25 to 4 in steps of 2^(1/4), giving 289 separate invasion analyses, identifying the ESS of each.

Based on the logic of Gadgil and Bossert (1970) and Schaffer (1974b) we predict that for the nine gain-loss treatments and the full sweep (i) our convex and linear treatments (*β* ≤ 1) should be invasible by a semelparous male at every other resident investment strategy (the PIPs should contain no interior ESS) and (ii) our concave treatments (*β* > 1) should include an interior investment strategy that is not invasible by any other strategy and (iii) that the full sweep analysis will reveal an invasion boundary for semelparous mutants at the linear treatments (*β* = 1) – i.e. semelparity will be the ESS for all cells with *β* ≤ 1 and iteroparity will be the ESS for all cells with *β* > 1. If these predictions do not hold, this means that features of our model that are absent from the original optimal allocation models – such as age-dependent survival and fecundity, density dependent survival or frequency dependent male paternity – are impacting the solution to the reproduction– survival trade-off.

### 2.6 – H2: Testing the predictions of the demographic model

To test the predictions of **the demographic model**, that higher and even lethal reproductive investment should be increasingly selected as post-reproductive survival declines, we systematically vary the timing of the onset of male post-reproductive survival decline. For the nine gain-loss treatments we projected our resident populations forward across a grid of male resident investment levels (*I_res_* ∈ {0.05, 0.10, …, 0.95}) and male mortality schedules at which monthly male survival reaches 50% (*m_fall_*_,*m*_ ∈ {12,13, …, 23} months), while holding female vital rates and gain and loss functions fixed, to determine which strategies are invasible by a semelparous mutant male. We then additionally conducted a full sweep analysis, sweeping a 9 × 9 grid of *γ*_*gain*_ and *γ_loss_*spanning 0.25 to 4 in steps of 2^(1/2) at each of the twelve onset ages, giving 972 separate invasion analyses and identifying the ESS of each.

We predict that if our model follows the logic of the demographic model then (i) for all three treatments where the reproduction-survival trade-off is convex (*β* < 1), semelparity will be the ESS regardless of the male mortality schedules, (ii) for all six of the gain-loss treatments where the trade-off is linear or concave (*β* ≥ 1), semelparity will be more likely to invade under male survival schedules with early post-reproductive survival declines – i.e. when males die earlier selection will favour semelparity – (iii) that for the full sweep analysis that the boundary separating iteroparous and semelparous treatments will monotonically move into increasingly concave regions of the trade-off plane as the onset becomes earlier and (iv) that treatments with more concave trade-off shapes will be more resilient to the semelparous mutant – i.e. the more concave the reproduction-survival trade-off, the earlier the male survival decline will need to be to make the population invasible.

### 2.7 – H3: Testing the predictions of the reproductive effort model

The **reproductive effort model** predicts that male-only semelparity can evolve when sexual competition selects for traits that trade off survival and future reproduction for current increased paternity and does not require post-reproductive mortality to favour semelparity (Fisher & Salguero-Gómez, 2026). To test the prediction, we alter our mating function to allow for high investing males to win a disproportionate proportion of paternity relative to their total mating supply. This allows us to capture the dynamics of sexual competition, where males compete over their share of limited number possible births by investing in traits like mate guarding or larger ejaculates. The strength of this skew is controlled by an exponential function with the parameter *φ* (Equation 8).

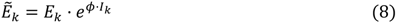

Where *k* is the male type, *E_k_* is its unweighted mating supply, *I_k_* is its reproductive effort and *EPk* is its mating supply after applying the paternity bias. When *φ* = 0 paternity is allocated between male types strictly proportionally to each type’s mating supply. However, when *φ* > 0, paternity is biased towards higher investing males i.e. they win more births than they should, given their reproductive investment and the shape of their underlying reproduction-survival trade-off curve.

To run an invasion analysis for each combination of gain and loss functions, we project our resident populations forward across a grid of possible male resident investment levels *I_res_* ∈ {0.05, 0.10, 0.15, …, 0.95}) and paternity skew strengths (*φ* ∈ {0, 0.5, 1, …, 3}) before introducing a rare semelparous mutant (*I_mut_* = 1). We then additionally conduct a full sweep analysis, sweeping a 9 × 9 grid of *γ*_*gain*_ and *γ_loss_* spanning 0.25 to 4 in steps of 2^(1/2)at each of the seven levels of paternity bias, giving 567 separate invasion analyses and identifying the ESS of each.

We predict that (i) across all nine combinations of gain and loss functions all investment strategies will be invasible for some value of *φ* leading to an identifiable invasion boundary, (ii) that the shape and location of this boundary will differ between combinations of gain and loss functions, and (iii) that for the full sweep analysis areas of the trade-off plane with higher *β* (more concave) will only be invasible at higher values of *φ*.

## 3. RESULTS

### 3.1 – H1: Invasion analysis support the predictions of the optimal allocation model for male-only semelparity

The classical optimal allocation model predicts that semelparity will evolve whenever increased reproductive investment has accelerating fecundity gains or decelerating survival costs. Our results demonstrate that in fact the shape of the reproduction-survival trade-off only makes semelparity possible but not inevitable because of its interaction with age-dependent vital rates. For all nine gain-loss treatments, the PIPs (Figure 3) identify a candidate ESS that cannot be invaded by any neighbouring strategy. For seven treatments this is a single strategy; for two of the three treatments with a linear trade-off (β = 1) it is a narrow band of adjacent strategies, none of which can be invaded (Table 2). Every candidate ESS was convergence stable. Residents investing less were invaded by higher-investing mutants and, for intermediate ESSs, residents investing more were invaded by lower-investing mutants. Selection therefore moves the population towards the ESS (Geritz et al., 1998). The ESS of female investment was *I_f_* = 0.3 for all nine treatments.

**Figure 3.**
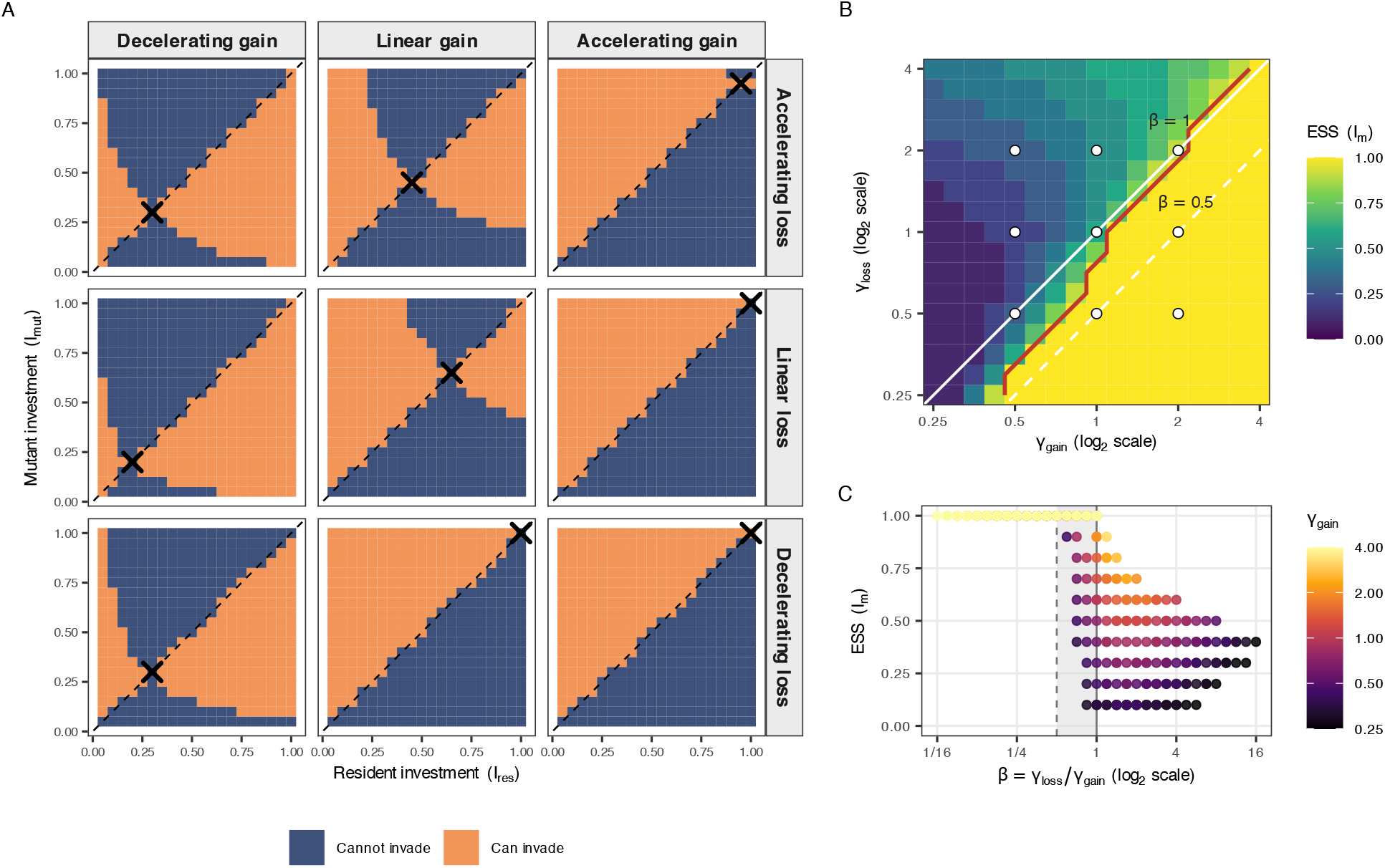
**(A)** Pairwise Invasibility Plot of reproductive investment strategies under each of our nine combinations of gain-loss functions. The x-axis shows the reproductive investment of the resident males *I_res_* and the y-axis the reproductive investment of the invading mutant male *I_mut_*. Orange squares indicate that at this combination of resident and mutant investment levels, the mutant population is growing relative to the resident population (*λ_inv_* > 1) and the resident strategy is invasible, while the blue squares show the regions where the resident population is declining and not invasible (*λ_inv_* < 1). The dashed line shows where the resident and mutant investment strategy are equal, and the black cross indicates the position of the Evolutionarily Stable Strategy (ESS), the resident investment strategy that cannot be invaded by any mutant investment strategy. For three of the treatments (accelerating gains ∼ decelerating losses, accelerating gains ∼ linear losses and linear gains ∼ decelerating losses) the ESS is semelparity (*I_res_* = 1). **(B)** Heatmap depicting the evolutionarily stable male reproductive investment across the reproduction-survival trade-off plane (17x17, n = 289). Each cell is a combination of *γ*_*gain*_ and *γ_loss_* values. For each value the uninvasible strategy was located by a full pairwise invasibility analysis as per section 2.4. The fill gives the ESS value of the gridcell and the red contour bounds the area beyond which semelparity (*I_m_* = 1) is the ESS. The axes are log2 scaled so that lines of constant *β* run at 45°, with the solid white line showing where *β* = 1 and the white dashed line where *β* = 0.5. The white circles mark the location of the nine treatments that comprise Figure 3A. **(C)** Each point is one of the 289 combinations of *γ*_*gain*_ and *γ_loss_* plotted against *β* and the ESS investment. The colour of the point encodes the *γ*_*gain*_ of the treatment combination. The grey area and the dashed line show the region 0.5 < *β* < 1, where treatments begin to progressively collapse towards semelparity.

**Table 2.**
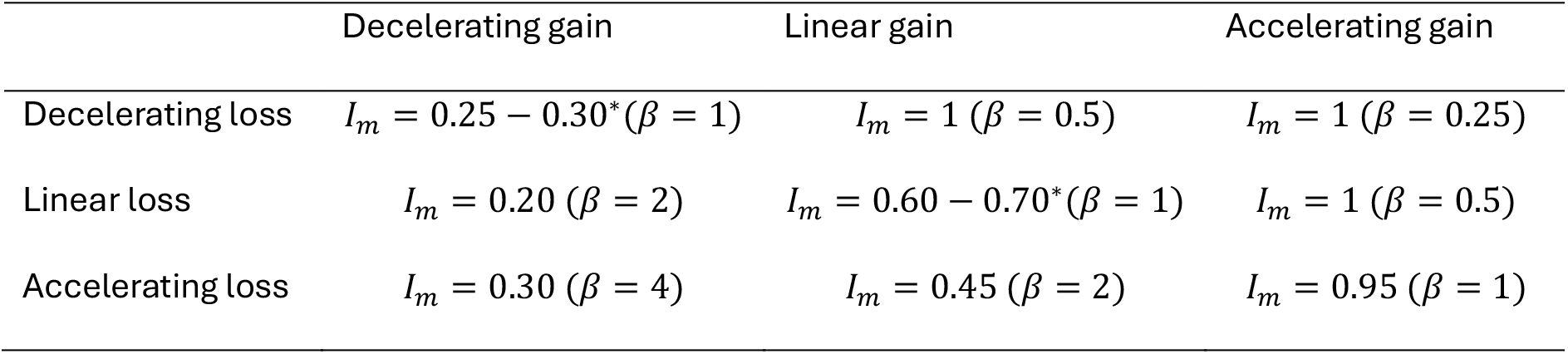
Evolutionary stable male reproductive investment strategy for each combination of gain function and loss function and the shape of their reproduction-survival trade-off (*β* = *γ_loss_*/*γ*_*gain*_). Several adjacent strategies (*) were uninvadable to within numerical precision (|*λ_inv_* − 1| < 10^–4^) so the ESS lies between them.

|  | Decelerating gain | Linear gain | Accelerating gain |
|---|---|---|---|
| Decelerating loss | *I*<sub>*m*</sub> = 0.25 − 0.30<sup>\*</sup>(*β* = 1) | *I*<sub>*m*</sub> = 1 (*β* = 0.5) | *I*<sub>*m*</sub> = 1 (*β* = 0.25) |
| Linear loss | *I*<sub>*m*</sub> = 0.20 (*β* = 2) | *I*<sub>*m*</sub> = 0.60 − 0.70<sup>\*</sup>(*β* = 1) | *I*<sub>*m*</sub> = 1 (*β* = 0.5) |
| Accelerating loss | *I*<sub>*m*</sub> = 0.30 (*β* = 4) | *I*<sub>*m*</sub> = 0.45 (*β* = 2) | *I*<sub>*m*</sub> = 0.95 (*β* = 1) |

Male-only semelparity (*I_m_* = 1) is the ESS for three of the nine gain-loss treatment combinations (Table 2). This includes the accelerating gains and decelerating losses, accelerating gains and linear losses, and linear gains and decelerating losses treatments. The PIP shows that under these treatments all resident strategies below semelparity (*I* < 1) are invasible by a higher investing mutant male (Figure 3A). This is consistent with the predictions of the Optimal Allocation Model, that semelparity will evolve when gains are accelerating or losses are decelerating (Gadgil & Bossert, 1970).

For all other treatment combinations, the ESS was an intermediate level of male reproductive effort (*I_m_* < 1). However, there is considerable variation in the ESS between these iteroparous treatments. When gains are accelerating (which under the optimal allocation model favours semelparity) but losses are also accelerating (which favours iteroparity), the ESS is near semelparity (*I_m_* = 0.95). In contrast, when gains and losses are decelerating, the ESS is much lower (*I_m_* = 0.25 − 0.30). This suggests that even though our gain and loss functions are symmetrical, their relative impacts on the model’s behaviour are not.

For the full sweep analysis semelparity was the male ESS in 89% of the treatments with a convex reproduction-survival trade-off (*β* < 1), 24% of the treatments with a linear trade-off (*β* = 1), and none of the treatments with a concave trade-off (*β* > 1). Semelparity becomes increasingly likely as the reproduction-survival trade-off becomes increasingly convex – semelparity was the ESS in 100% of treatments with *β* < 0.5. Among those treatments where the ESS was iteroparity, *β* explained almost none of the variation in the actual optimal amount of male reproductive investment *R*^2^ = 0.078, against *R*^2^ = 0.704 for log *γ*_*gain*_ and *R*^2^ = 0.178 for log *γ_loss_*. This suggests that while *β* ≤ 1 may be the criterion for the evolution of semelparity, among the iteroparous treatments, the shape of the gain function plays a more important role in determining the optimal level of reproductive investment compared to the loss function.

Changing the *b_dd_* in our sensitivity analysis altered neither the ESS nor the invasion outcome, showing that the differences between treatments were driven by the shape of the trade-off geometry itself not its interaction with the density-dependence term (Supporting Information S5, Table S1).

### 3.2 – H2: The demographic model is constrained by the underlying reproduction-survival trade-off

As the onset of post-reproductive male mortality becomes earlier – so the post reproduction survival rate declines – the capacity of the semelparous mutant to invade increased under every gain-loss treatment (Figure 4A), but invaded a stable iteroparous population only when the trade-off was close to or on the semelparity threshold. Higher investment strategies of residents generally buffered against the semelparous mutant invading. The three treatments with a semelparous ESS were invasible by a semelparous mutant for all values of *m_fall_* and *I_res_*. The accelerating gain ∼ accelerating loss treatment (near-semelparous ESS), was invasible at all mortality onsets except at the highest resident investment. All three treatments with decelerating gain functions resisted the semelparous mutant, which invaded only when post-reproductive survival declined extremely early (12-15 months) and resident investment was low. Higher post-reproductive male mortality can therefore create the conditions for the evolution of semelparity. However, the exact age at which that threshold is crossed is depends on both the shape of the reproduction-survival trade-off and its interaction with existing reproductive investment.

**Figure 4.**
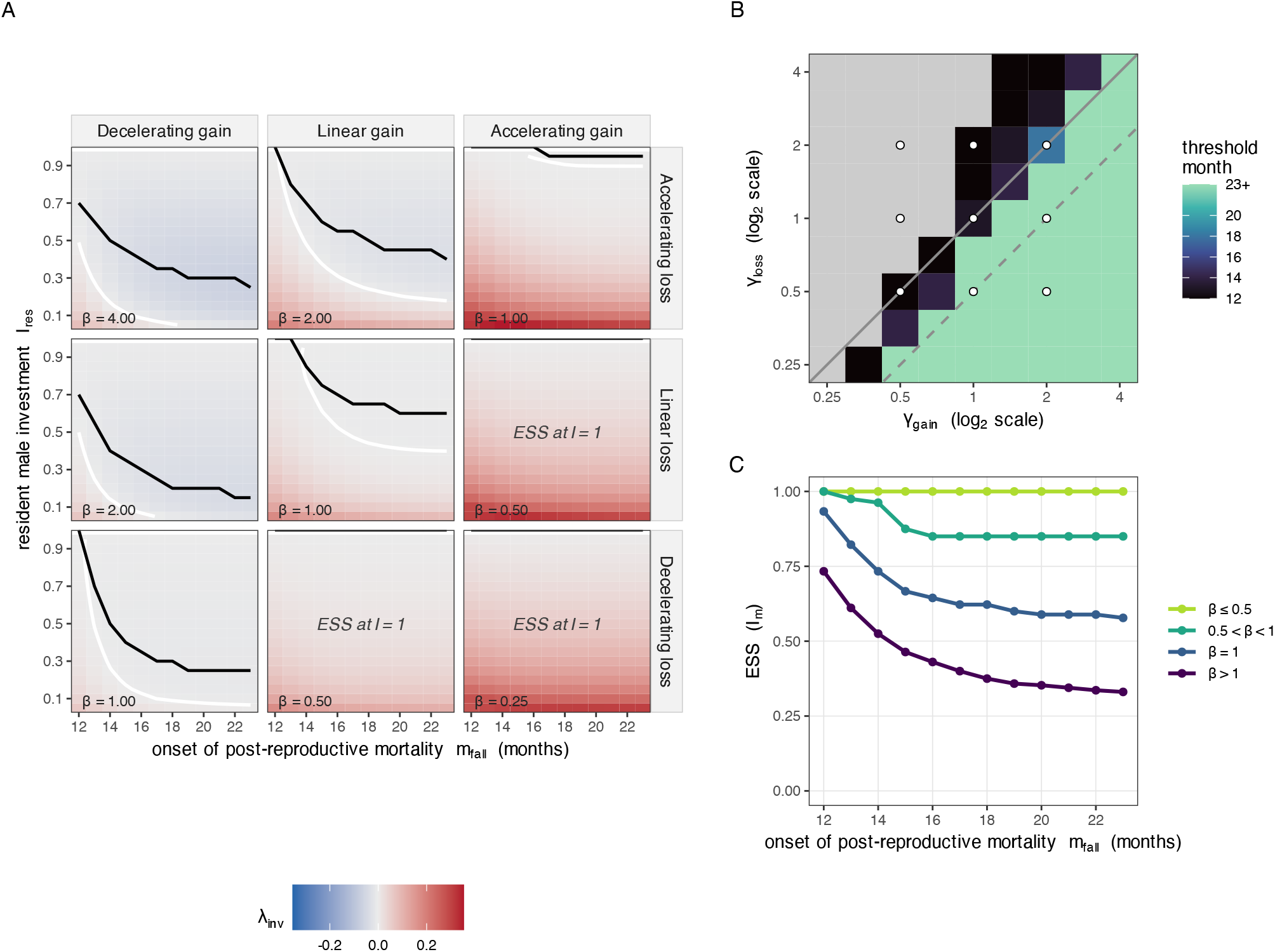
**(A)** Invasion surface depicting the invasibility of a semelparous mutant male across different male resident investment (*I_res_*) and post-reproductive mortality schedules (*m_fall_*). The colour of the cell indicates the geometric per time step growth rate of the invading semelparous mutant (*λ_inv_*). If *λ_inv_* > 1 then the semelparous male abundance is growing relative to the resident males and is invading the population. The white line shows the invasion boundary that separates the region the mutant male can invade from the region it cannot invade, while the black line shows the male investment that maximises population growth rate for each post-reproductive mortality schedule, which matches the evolutionarily stable investment from the invasion analysis to within the grid resolution (±0.05; Supplementary Data SD4). **(B)** Each cell is one combination of gain and loss function shape, with *γ*_*gain*_ and *γ_loss_*. The fill gives the latest month from 12 to 23 months where semelparity (*I_m_* = 1) is the ESS. Cells with 23+ are always semelparous and grey cells are never semelparous, regardless of the age at which post-reproduction survival decline begins. Axes are log₂-scaled so that lines of constant *β* = *γ_loss_*/*γ*_*gain*_run at 45°: the solid diagonal is *β* = 1and the dashed diagonal is *β* = 0.5. Circles mark the nine treatments of Figure 4A. **(C)** Coloured lines show the mean of the evolutionarily stable male investment *I_m_* within four bands of *β* as the onset of post-reproductive mortality varies from 12 to 23 months. Earlier onset of post-reproduction mortality selects for increased male investment but only when the shape of the reproduction-survival trade-off permits it. The effect is absent when *β* ≤ 0.5 where semelparity is the ESS. Above this threshold, most of the change occurs before the age of 17 months, after which additional post-breeding survival has little effect.

An earlier onset of post-reproductive mortality never decreased male reproductive investment. For the full sweep analysis the onset of post-reproductive mortality moved the ESS in 46 of the 81 cells tested and the mean evolutionarily stable level of male reproductive investment rose from *I_m_* = 0.64 when post-reproductive survival declined at 23 months to *I_m_* = 0.87 when it declined at 12 months(Figure 4B, Figure 4C). This movement was not confined to cells with linear or near-linear trade-offs. Every cell with a concave reproduction-survival trade-off (*β* > 1) responded to changes in the mortality schedule (36 of 36 cells), and by the largest margin of any band, with ESS *I_m_*increasing on average by 0.40 as the onset of mortality moved from 23 to 12 months. Most of the linear cells (*β* = 1) responded as well (7 of 9 cells), with a mean increase of 0.36, while cells with 0.5 < *β* < 1 were the least responsive of all (3 of 8 cells; mean increase 0.15). The only regions which did not respond were those with very convex reproduction-survival trade-off curves (*β* ≤ 0.5) where semelparity was the ESS.

Responsiveness to the mortality schedule and the capacity to reach semelparity are, however, distinct properties and our results suggest that the demographic model can only lead to the evolution of a semelparous population under some reproduction-survival trade-off shapes. For example, the concave cells in the full sweep analysis moved the most while arriving at *I_m_* = 1 the least often, with only 8 of 36 ever doing so. In total only 16 cells flipped from iteroparity to semelparity as the onset of post-reproductive mortality became earlier, spanning *β* = 0.71, 1, 1.41, 2 and 2.83 (Figure 4B). The capacity to flip from iteroparity to semelparity therefore reaches well into the concave region of the trade-off surface but only seems to be feasible when either the age of onset of post reproduction mortality is very early (11 of 16 cells flipped at 12 to 13 months, just one to two months after males reach sexual maturity) or only when the reproduction-survival trade-off shape was *β* ≤ 2.83. None of the 15 cells with extremely concave trade-offs *β* ≥ 4 ever flipped from iteroparity to semelparity. This suggests that for our model, early onset of post-reproductive mortality can select for higher male reproductive investment across the whole trade-off plane and can drive linear and some concave trade-offs all the way to semelparity consistent with the predictions of age-structured demographic models. However, this is mostly only true when post-reproduction mortality begins within a few months of sexual maturity and not when the reproduction survival trade-off is strongly concave.

### 3.3 – H3: Supporting the reproductive effort model: sexual competition can lead to semelparity

Under every gain-loss treatment, some level of paternity skew (*φ*) allowed a semelparous mutant to invade all iteroparous resident male investment strategies (*I_res_* < 1) but the skew varied considerably between treatments (Figure 5A). As in H2, the three treatments with a semelparous ESS were invasible for any combination of *I_res_* and *φ*. For the decelerating gain ∼ decelerating loss treatment *φ* = 0.5 made all *I_res_* < 1 strategies invasible. Under this treatment if resident and mutant males contribute equal mating supply, so that without paternity skew each would sire half the births, a resident strategy of *I_res_* = 0.95 and paternity skew of *φ* = 0.5 give the semelparous mutant 50.62% of births (Figure 5B). This 0.62 percentage point advantage is enough for it to invade. Under decelerating gains and accelerating losses, a resident with *I_res_* = 0.40 could only be invaded when *φ* > 1. Again, assuming equal proportions of semelparous and resident male mating supply in the population, if *φ* = 1 and *I_res_* = 0.40 the semelparous invaders are winning 64.57% of the births and cannot invade. Female investment was held at *I_f_* = 0.3, because *φ* only redistributes paternity among males, it does not alter selection on females. Semelparity can therefore evolve through sexual competition alone if reproductive investment translates to a disproportionate share of paternity, but the skew required rises steeply as the underlying trade-off geometry increasingly favours iteroparity.

**Figure 5.**
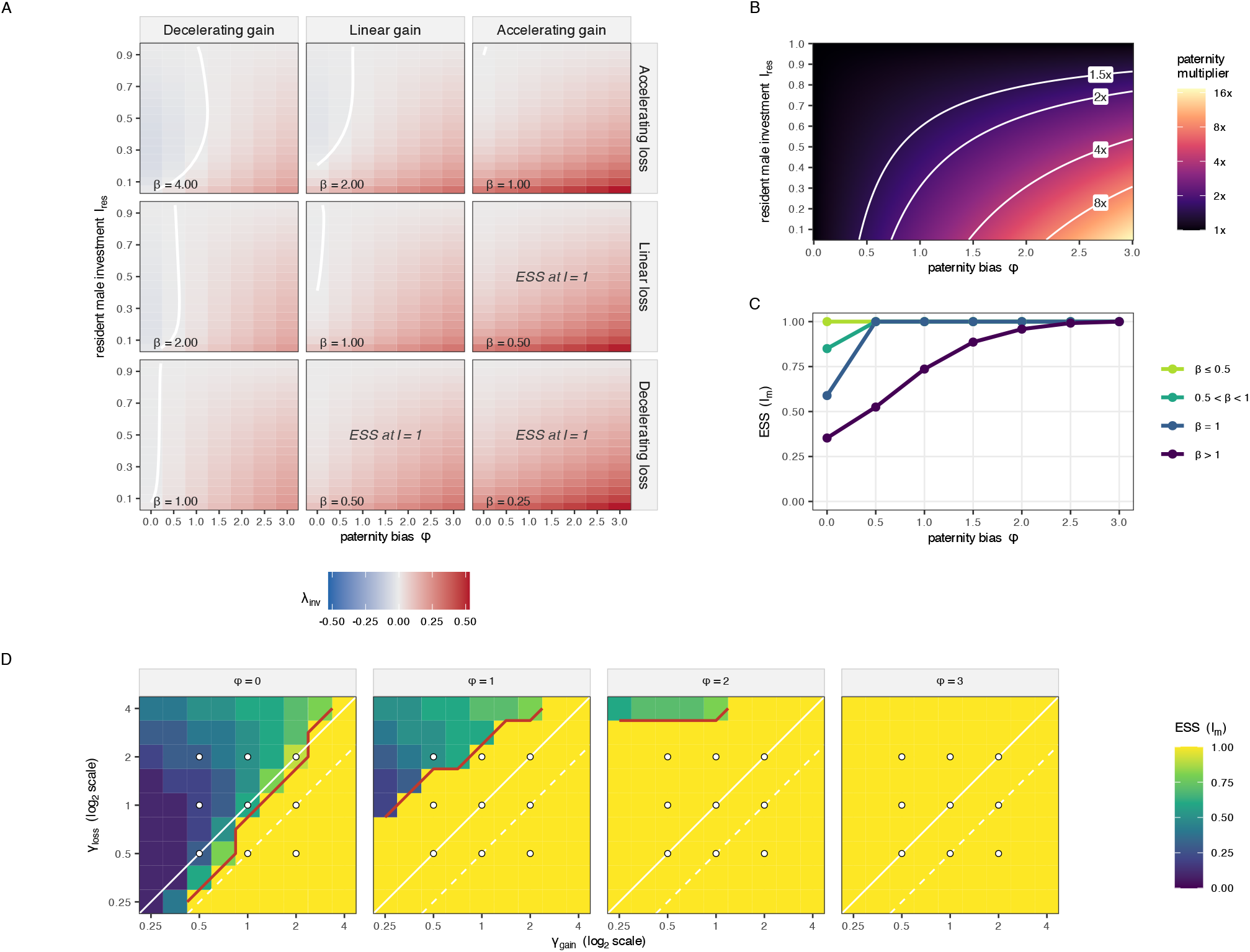
**(A)** Invasion surface depicting the invasibility of a semelparous mutant male across different male resident investment (*I_res_* = 0.05 − 0.95) and paternity bias (*φ* = 0 − 3) for each of the nine male trade-off treatments. The colour of the cell indicates the geometric per timestep growth rate of the invading semelparous mutant (*λ_inv_*). If *λ_inv_* > 1 then the semelparous male abundance is growing relative to the resident males and is invading the population. The white contour shows the invasion boundary that separates the region the mutant male can invade from the region it cannot invade. **(B)** The factor by which paternity bias multiplies a semelparous mutant’s (*I_mut_* = 1) share of paternity relative to allocation by mating supply alone across paternity bias (*φ* = 0 − 3) and male resident investment (*I_res_* = 0.05 − 0.95). White contours give the locations of 1.5x, 2x, 4x and 8x multipliers. Colours are on a log2 scale. **(C)** Evolutionarily stable male investment against *φ*, averaged within four bands of *β* across the full 9x9 reproduction-survival trade-off sweep. When there is no paternity bias towards higher investing males (*φ* = 0) the geometry of the trade-off splits the ESS. However, as *φ* increases, it selects for increasingly high and even lethal male investment. By *φ* = 1 all but the concave trade-off band are driven to semelparity. **(D)** ESS male investment across the 9x9 reproduction-survival trade-off sweep at *φ* = 0, 1, 2 and 3. Both the x-axis (*γ*_*gain*_) and y-axis (*γ_loss_*) are log2-scaled so that the lines of constant *β* are 45° diagonals (solid white, *β* = 1; dashed white, *β* = 0.5). The red contour separates the semelparous region of the landscape from the iteroparous region, and the white circles mark the location of the nine treatments. The semelparous region expands as *φ* increases, driving linear and near linear regions to semelparity first. The areas with a concave trade-off require much higher values of *φ* for semelparity to be the ESS.

For the full sweep analysis when *φ* = 0, where paternity is allocated strictly in proportion to mating supply, the landscape was consistent with that of section 3.1. Increasing the paternity bias expands the region of the trade-off surface where semelparity is the ESS – 51 cells when *φ* = 0.5, 62 at *φ* = 1, 76 at *φ* = 2, and the entire grid at *φ* = 3 (Figure 5C, 5D). Unsurprisingly, cells with linear or convex trade-offs were the most sensitive to paternity bias – all cells with *β* < 1 were semelparous at *φ* = 0.5. In contrast to the onset of post-reproductive mortality, where trade-offs that were more concave than *β* = 2.83 never flipped from iteroparity to semelparity, a sufficiently high paternity bias of *φ* = 3 was able to make the ESS semelparity for all cells.

Unlike the age-structured demographic model (section 3.2), paternity bias changed the degree to which different components of the trade-off geometry affected the ESS of iteroparous treatments. log_2_ *γ*_*gain*_ was the strongest single predictor at every level of *φ* (R^2^= 0.72-0.86), but the effect of the loss function more than tripled between *φ* = 0 and *φ* = 1 (R^2^= 0.20-0.66). Paternity biases amplify fecundity returns of higher investment, such that survival costs rather than fecundity gains become the limiting factor on where an interior ESS can sit. Although, we note that these values should be interpreted with caution, as the number of treatments with an interior ESS declined from 46 at *φ* = 0 to only five at *φ* = 2, with the remaining treatments being largely concave.

Paternity skew is sufficient to make semelparity the male ESS. If paternity is a strictly zero-sum game and if high-investing males receive a share of paternity disproportionate to their investment and at the expense of low-investing males, semelparity can invade populations that would otherwise be iteroparous while leaving females unchanged. However, there is an important caveat to this interpretation. Because the fecundity returns of investing in sexual competition traits like sperm competition or mate guarding are unknown in *A. agilis,* whereas the onset of post-reproductive mortality is bounded by demography data for this species (section 3.2), we cannot say which mechanism is stronger, only that both are viable paths to semelparity over some portion of the trade-off plane.

The main findings for each test are summarised in Table 3.

**Table 3.** Summary of main findings.

|  | **H1**<br>Optimal allocation model | **H2**<br>Age-structured demographic model | **H3**<br>Reproductive effort model |
|---|---|---|---|
| *What is tested* | Male ESS of reproductive investment across different γ<sub>gain</sub> and γ<sub>loss</sub> combinations | Impact of the onset of post-reproductive mortality (m<sub>fall</sub> = {12-23 months}) on male ESS | Impact of paternity bias (ϕ = {0-3}) on the male ESS |
| *Main findings* | A convex or linear trade-off β ≤ 1 is necessary for semelparity but not sufficient.<br>The γ<sub>gain</sub> determines the position of internal ESS. | Early onset of post reproductive male mortality leads to increased reproductive investment and in some cases can flip a population from iteroparity to semelparity but never the reverse.<br>The γ<sub>gain</sub> determines the position of internal ESS across all onsets of mortality. | Paternity share bias towards high-investing males allows semelparity to invade reproduction-survival trade-offs that would otherwise be iteroparous. Increased paternity bias increases the role of γ<sub>loss</sub> in determining internal ESS. |
| *Do the findings support the model's predictions* | Yes, but the shape of the reproduction-survival trade-off makes semelparity *possible* not *inevitable* because of its interaction with age-dependent vital rates. | Yes, but only when the trade-off geometry is close to or on the threshold for semelparity threshold, and mostly only when mortality begins shortly after sexual maturity. | Yes, paternity bias can potentially lead to semelparity in all regions of the reproduction-survival trade-off plane. However, careful consideration should be given to what is a realistic level of paternity bias for a given investment strategy. |
| *Reconciling findings with the life-history of species with male-only semelparity* | For male-only semelparity to evolve, males and females must have either (i) different shaped reproduction-survival trade-offs or (ii) similar but borderline trade-offs, with different vital rates(→H2). | It could be the difference in demography explains male-only semelparity under H1 but requires some factor extrinsic to reproduction that makes mortality much severe in post reproductive males than females. | Reproductive strategies that reward high investment through biased paternity (e.g. mate guarding, sperm competition) could lead to male-only semelparity. However, many metazoans exhibit such traits and yet few exhibit male-only semelparity.<br>Implies the existence of a constraint that male-only semelparous species have escaped. |

## 4. DISCUSSION

In animals, a semelparous life-history is as rare as it is extreme, and theoretical work into understanding the conditions under which it can evolve emphasise either the geometry of the underlying reproduction-survival trade-off (optimal allocation model) or the way extrinsic mortality risk changes over an organisms lifetime, particularly after reproduction (age-structured demographic model). However, it has not been clear how these models apply to species with male-only semelparity. Indicators of intense sexual competition among males in these species suggest that sexual competition may itself be a pathway to the evolution of semelparity (the sexual competition model). Using invasion analysis of a two-sex, age-structured population model, we found that all three pathways allow semelparity to become the evolutionarily stable strategy in males, even when females remain iteroparous. However, each model leads to semelparity in different and largely non-overlapping regions of the trade-off plane. These are not mutually exclusive hypotheses, but separate, potentially interacting evolutionary forces that can make male-only semelparity a viable life-history strategy. The geometry of the reproduction– survival trade-off sets the threshold at which semelparity becomes accessible, while demography and sexual competition shift that threshold, determining whether semelparity evolves.

Our test of the optimal allocation model shows that a convex reproduction-survival trade-off is a criterion for the evolution of semelparity but does not make semelparity inevitable. Iteroparity will always evolve when the trade-off is concave, but it is also evolutionarily stable in some linear and convex regions of the trade-off plane, where the classical model would predict semelparity (Gadgil & Bossert, 1970; Schaffer, 1974a), suggesting that our age-structured vital rates and density dependent population growth are maintaining iteroparity in these regions (Benton & Grant, 1999; Charnov & Schaffer, 1973; Schaffer, 1974b). A species’ reproduction-survival trade-off curve can rarely be measured in the field (Reznick et al., 2026; van Noordwijk & de Jong, 1986), although experimental and comparative studies have described its shape or tested its predictions in some systems (Fisher, Double, Blomberg, et al., 2006; Gustafsson & Sutherland, 1988; Kindsvater et al., 2016; Santos & Nakagawa, 2012). But a reproduction-survival trade-off is a simplification of the interactions among all aspects of an organism’s life-history including its demography? Rather than capturing processes separate from the demographic and reproductive effort models, the optimal allocation model should be understood as the background against which they act.

Both the demographic model and reproductive effort model raise male reproductive investment, but in markedly different ways. Earlier onset of post-reproduction mortality was associated with greater male reproductive investment and in some circumstances converted iteroparous populations to semelparity. This is consistent with several theoretical studies that report that increased juvenile survival relative to adult survival, earlier ages of senescence and decreased age of sexual maturity favour semelparity (Bulmer, 1985; Charnov & Schaffer, 1973; Young, 1981). Unexpectedly, the most concave regions of the reproduction-survival plane, and particularly those with decelerating gain functions, proved resistant to the effects of an earlier onset of post-reproduction mortality. The capacity of the demographic model to shift a population from iteroparity to semelparity is constrained by the underlying geometry of the reproduction-survival trade-off.

For the demographic model to act as a pathway to semelparity an aspect of an organism’s biology or environment must increase post-reproductive mortality. For example, in aquatic animals semelparity is typically associated with costly migrations to breeding grounds at sexual maturity (Crespi & Teo, 2002; Fisher & Salguero-Gómez, 2026; Kindsvater et al., 2016). The cause of high post-reproductive mortality in semelparous terrestrial animals (and plants) is typically seasonally harsh (cold, hot, or arid) environments in which suitable conditions for reproduction are punctuated by conditions that greatly increase mortality risk (Fisher & Salguero-Gómez, 2026; Guillet & Vancassel, 2001; Karsten et al., 2008; Smith & Charnov, 2001; Young, 1990). We know of only one example in which an aspect of an organism’s biology or environment unambiguously increases post-reproductive mortality in only one sex, associated with semelparity in that sex and iteroparity in the other sex. European conger eels (*Conger conger*), appear to be the only known species to exhibit female-only semelparity, and it is hypothesised that this is a consequence of sex difference in spawning migration. Only females migrate from shallow waters to deep ocean regions, where the males are permanent residents (Casadevall et al., 2017). Both sexes of *Anguilla* eels experience long, demanding spawning migrations to locations in the deep ocean and both sexes are semelparous (Jellyman, 2022; Tsukamoto et al., 2011).

Our model shows that sexual competition among males for paternity can select for male-only semelparity, consistent with experimental and comparative data in dasyurids and spiders (Andrade, 1996; Dickman, 1994; Fisher et al., 2013; Forster, 1992). When paternity-skew towards high-investing males was sufficiently high, male post-mating competition was able to change the entire trade-off landscape to semelparity. This is an abstraction of the sexual competition traits hypothesised to select for male-only semelparity, such as sperm competition (Dickman, 1994; Fisher et al., 2013) and mate guarding (Andrade, 1996; Forster, 1992). Our model does not include female preferences, female-mediated competition or coevolution between the sexes, and so captures competition among males rather than sexual conflict. Increasing the fitness gain for high-investing strategies of males shifts the semelparity threshold into more concave areas of the trade-off plane that do not maximise overall population growth. From the perspective of male fitness, adopting a strategy that allows males to dominate paternity, even if it leads to fewer births in the population overall, is a fitter strategy (Kokko & Brooks, 2003; Rankin et al., 2011). However, it is difficult to assess whether the paternity returns required for semelparity to be a viable evolutionary strategy in the most concave regions of the trade-off plane are biologically plausible. Holleley et al. (2006) found that paternity success was most strongly related to body mass and scrotal size in *Antechinus stuartii,* suggesting that in this male-only semelparous species increased reproductive investment does translate into increased paternity success.

Studies of individual species including mantids (Brown & Barry, 2016; Hurd, 1989) and hymenoptera (Baer, 2005), and comparative tests of dasyurids (Fisher et al., 2013) and spiders (Miller, 2007; Schwartz et al., 2013, 2014), have demonstrated close associations between male-only semelparity and traits that indicate intense post-copulatory sexual competition. In dasyurids these traits are associated with mate-guarding to protect paternity: exceptionally prolonged copulations, large testes, and physiological die-off syndrome that depletes body energy and nitrogen enabling concentrated effort, endurance, and wakefulness (Fisher et al., 2013; Fisher & Salguero-Gómez, 2026; Lazenby-Cohen & Cockburn, 1988). In arthropods, traits include other extreme forms of mate-guarding to protect paternity: semelparous male spiders lethally fasting while mate-guarding (Cohn et al., 1988),lethally transferring body fluids to inflate the end of their pedipalp and block the female opening after death (Knoflach & Harten, 2001; Ramos et al., 2004; Schwartz et al., 2014), spiders, ants, and bees lethally amputating body parts to block the female opening after death(Baer, 2005; Kuntner et al., 2009; Monnin & Peeters, 1998),and males inciting sexual cannibalism that increases male fitness (Andrade, 1996; Brown & Barry, 2016).

Mate guarding and sperm competition are widespread (Fitzpatrick, 2020; Møller & Briskie, 1995; Soulsbury, 2010). If sexual competition selects for extreme male reproductive investment, why are there so few species with male-only semelparity? In the vast majority of animals that mate-guard, male effort has not escalated to the extent that males invest lethally in reproduction. Fisher, Double, Blomberg, et al. (2006) experimentally demonstrated that female brown antechinus gain substantial lifetime fitness benefits from promiscuity because they obtain good genes for their offspring through a mechanism of sperm competition, suggesting that selection favours females that escalate the intensity of male post-mating competition. Good genes mechanisms are rarely demonstrated as the explanation for the widespread phenomenon of female promiscuity (Fisher, Double, Blomberg, et al., 2006). It may be that selection for lethal mate-guarding requires a good genes mechanism, so selection can act on female incitement of male competition.

Conversely, sexual conflict could itself constrain male reproductive investment. Our model assumes that male and female reproductive investment are decoupled, but if they were linked, for example through a shared genetic basis, selection for increased male reproductive investment could be constrained by its costs to females. Intralocus sexual conflicts are widespread (Berg & Maklakov, 2012; Foerster et al., 2007; Mills et al., 2012), so it is possible that male-only semelparity can only evolve if the genetic architecture underpinning reproductive investment is decoupled between the sexes. A future study could extend our model to incorporate sexual conflict and ask whether treating male and female reproductive investment as non-independent traits affects the evolutionary viability of male-only semelparity.

Alternatively, although they are iteroparous, 85-90% of female agile antechinus die before weaning a second litter (Fisher & Blomberg, 2011). It may be that by basing our model on demography of female *A. agilis*, our vital rates are already close to semelparity, and so only slight paternity skew is needed to shift the landscape towards semelparity. For example, our model’s rate of juvenile survival is relatively high for a mammal (Oli & Dobson, 2003), and low juvenile survival can maintain iteroparity in some models (Bulmer, 1985; Charlesworth, 1994). Repeating these analyses with different baseline vital rates based on the demography of semelparous arthropods and other dasyurids would be instructive to test generality of these pathways.

Under our model, vital rates only changed in relation to the density-dependent term between time steps – we effectively assumed that the environment was stable. Environmental variation and (Bulmer, 1985; Murphy, 1968; Orzack & Tuljapurkar, 1989; Schaffer, 1974a) and predictability (Collett et al., 2018; Guillet & Vancassel, 2001) are important in shaping trade-offs between reproduction-and survival and maintaining iteroparity. In a year with no recruitment a one-year-old iteroparous male potentially faces a cohort with no rivals, an advantage that could push back against the advantages of biased paternity. A future study could extend our model to test the effects of environmental variation, looking at how the severity and predictability of that variation affects recruitment, and asking whether unpredictable recruitment failure can maintain iteroparity.

Published tests suggest that post-mating sexual selection under sexual conflict (in which females benefit from promoting costly competition among males) is the driver of male-only semelparity in animals (Fisher & Salguero-Gómez, 2026). Indicators of pre-mating sexual selection such as male ornaments and armaments are not associated with male-only semelparity (Fisher & Salguero-Gómez, 2026). Our model allows males to compete over paternity by translating reproductive effort into one immediate fecundity bonus, consistent with characteristics of post-mating sexual competition in semelparous male dasyurids and arthropods; sperm competition (or sperm selection by females) with suicidal mate guarding during or immediately following copulation, in which males invest their body tissue in defending paternity. This immediate benefit and cost may be a reason why post-mating (not pre-mating) competition is the driver of male-only semelparity. Pre-mating sexual selection such as ornaments to attract females, armaments to fight off rival males, and territories to secure exclusive access to females may be costly to grow or establish, but may persist in repeated seasons (Champagne et al., 2012; Karman et al., 2026; Lane et al., 2010 see table 2; Soulsbury, 2010). Iteroparity could be maintained, even when the underlying geometry favours semelparity, if males invest in a sexually selected trait that benefits them in repeated seasons. Another reason why post-mating (not pre-mating) competition is the driver of male-only semelparity is likely to be that if they have not yet mated, males cannot increase paternity by lethally high effort. However, several mechanisms increase paternity through death once mating has started or is completed, so post-mating investment in paternity at the expense of survival can be selected (Fisher & Salguero-Gómez, 2026).

## 5. CONCLUSION

All three of our proposed pathways can explain the evolution of male semelparity, and each is largely in a non-overlapping region of the reproduction-survival trade-off plane. While the criterion of the classical optimal allocation holds, that semelparity can evolve when the trade-off is either linear or convex, it sets the threshold at which semelparity is evolutionarily accessible while the demographic and sexual competition models shift that threshold. The onset of post-reproductive mortality can drive much greater levels of male reproductive investment but in our model, it can only convert iteroparous populations to semelparity when the underlying trade-off geometry is linear or near linear. Post-mating sexual competition is the only pathway that could produce semelparous populations across the entire trade-off plane, and it did so without any change to extrinsic mortality, lending support to the reproductive effort model. Yet it is difficult to ascertain what is a biologically plausible paternity skew for investing in sexual competition and the fact that male-only semelparity is rare in animals while sexual competition between males is extremely common, suggests that demography, environment or the peculiarities of mating systems act as a constraint on the evolution of lethal reproductive investment.

## Data Availability

Upon publication all code and data to replicate analyses will be made available on the Dryad Digital Repository.

## Author Contributions

**KJY**: Conceptualization (equal), formal analysis, investigation, software, methodology (lead), visualization, writing – original draft, writing – review & editing (lead). **RSG**: Conceptualization (equal), methodology (supporting), writing – review & editing (supporting), supervision (equal). **DOF**: Conceptualization (equal), methodology (supporting), writing – review & editing (supporting), supervision (equal).

## Acknowledgements

We acknowledge the Ngunnawal, Jagera and Turrbal people, the traditional custodians of the lands on which this manuscript was prepared, land which was never ceded, and we pay our respects to their Elders, past, present and emerging. We also thank Hanna Kokko, Claudia Crowther, Chrissy Hernandez and Sean Pang for their thoughtful suggestions and encouragement.

## REFERENCES

Ancona, S., Liker, A., Carmona-Isunza, M. C., & Székely, T. (2020). Sex differences in age-to-maturation relate to sexual selection and adult sex ratios in birds. Evolution Letters, 4(1), 44–53. 10.1002/evl3.156

Andrade, M. C. B. (1996). Sexual Selection for Male Sacrifice in the Australian Redback Spider. Science, 271(5245), 70–72. 10.1126/science.271.5245.70

Baer, B. (2005). Sexual selection in Apis bees. Apidologie, 36(2), 187–200. 10.1051/apido:2005013

Benton, T. G., & Grant, A. (1999). Elasticity analysis as an important tool in evolutionary and population ecology. Trends in Ecology and Evolution, 14(12), 467–471. 10.1016/S0169-5347(99)01724-3

Berg, E. C., & Maklakov, A. A. (2012). Sexes suffer from suboptimal lifespan because of genetic conflict in a seed beetle. Proceedings of the Royal Society B: Biological Sciences, 279(1745), 4296–4302. 10.1098/rspb.2012.1345

Bonduriansky, R., Maklakov, A., Zajitschek, F., & Brooks, R. (2008). Sexual selection, sexual conflict and the evolution of ageing and life span. Functional Ecology, 22(3), 443–453. 10.1111/j.1365-2435.2008.01417.x

Braithwaite, R. W., & Lee, A. K. (1979). A Mammalian Example of Semelparity. The American Naturalist, 113(1), 151–155. 10.1086/283372

Brooks, R. C., & Garratt, M. G. (2017). Life history evolution, reproduction, and the origins of sex-dependent aging and longevity. Annals of the New York Academy of Sciences, 1389(1), 92–107. 10.1111/nyas.13302

Brown, W. D., & Barry, K. L. (2016). Sexual cannibalism increases male material investment in offspring: Quantifying terminal reproductive effort in a praying mantis. Proceedings of the Royal Society B: Biological Sciences, 283(1833), 20160656. 10.1098/rspb.2016.0656

Bulmer, M. G. (1985). Selection for Iteroparity in a Variable Environment. The American Naturalist, 126(1), 63–71.

Casadevall, M., Sarra-Alarcon, L., Delgado, E., & Matallanas, J. (2017). The sexual segregation of the European eel, *Conger conger* (Linnaeus, 1758) (Anguilliformes, Congridae) and female semelparity in the northwest Mediterranean. Journal of Fisheries Research, 1, 5–14.

Champagne, C. D., Houser, D. S., Fowler, M. A., Costa, D. P., & Crocker, D. E. (2012). Gluconeogenesis is associated with high rates of tricarboxylic acid and pyruvate cycling in fasting northern elephant seals. *American Journal of Physiology – Regulatory*, Integrative and Comparative Physiology, 303(3), R340–R352. 10.1152/ajpregu.00042.2012

Charlesworth, B. (1994). Evolution in Age-Structured Populations (2nd ed.). Cambridge University Press.

Charnov, E., & Schaffer, W. (1973). Life-History Consequences of Natural Selection: Cole’s Result Revisited. The American Naturalist, 107(958), 791–793. 10.1086/282877

Cockburn, A., Scott, M. P., & Scotts, D. J. (1985). Inbreeding avoidance and male-biased natal dispersal in Antechinus spp. (Marsupialia: Dasyuridae). Animal Behaviour, 33(3), 908–915. 10.1016/S0003-3472(85)80025-7

Cohn, J., Balding, F. V., & Christenson, T. E. (1988). In defense of Nephila clavipes: Postmate guarding by the male golden orb-weaving spider. Journal of Comparative Psychology, 102(4), 319–325. 10.1037/0735-7036.102.4.319

Cole, L. C. (1954). The Population Consequences of Life History Phenomena. The Quarterly Review of Biology, 29(2), 103–137.

Collett, R. A., Baker, A. M., & Fisher, D. O. (2018). Prey productivity and predictability drive different axes of life-history variation in carnivorous marsupials. Proceedings of the Royal Society B: Biological Sciences, 285(1890), 20181291. 10.1098/rspb.2018.1291

Crespi, B. J., & Teo, R. (2002). Comparative phylogenetic analysis of the evolution of semelparity and life history in salmonid fishes. Evolution, 56(5), 1008–1020. 10.1111/j.0014-3820.2002.tb01412.x

Cushing, J. M. (2021). A bifurcation theorem for Darwinian matrix models and an application to the evolution of reproductive life-history strategies. Journal of Biological Dynamics, 15(sup1), S190–S213. 10.1080/17513758.2020.1858196

Dickman, C. R. (1989). Demographic responses of *Antechinus stuartii* (Marsupialia) to supplementary food. Australian Journal of Ecology, 14(4), 387–398. 10.1111/j.1442-9993.1989.tb01449.x

Dickman, C. R. (1994). Evolution of semelparity in male dasyurid marsupials: A critique, and an hypothesis of sperm competition. In J. Carnio (Ed.), *Carnivorous Marsupials* (pp. 19–32). Metro Toronto Zoo.

Fairbairn, D. J. (1997). Allometry for Sexual Size Dimorphism: Pattern and Process in the Coevolution of Body Size in Males and Females. Annual Review of Ecology and Systematics, 28(1), 659–687. 10.1146/annurev.ecolsys.28.1.659

Ferreira, M. S., Rossi, R. C., Batista, S., Sena, K., Bairos-Novak, K. R., Shoma, S. F., Leiner, N. O., Astúa, D., Coulson, G., Fisher, D. O., Harrison, N. D., Willmer, J. N. G., Silva, G. R., Cupolillo, G., Vieira, M. V., Vieira, E. M., Püttker, T., Figueiredo, M. de S. L., & Dickman, C. R. (2026). The Marsupial Database: A comprehensive dataset on the ecology and life history of American and Australasian marsupials. Ecology, 107(3), e70358. 10.1002/ecy.70358

Fisher, D. O., & Blomberg, S. P. (2011). Costs of Reproduction and Terminal Investment by Females in a Semelparous Marsupial. PLOS ONE, 6(1), e15226. 10.1371/journal.pone.0015226

Fisher, D. O., Dickman, C. R., Jones, M. E., & Blomberg, S. P. (2013). Sperm competition drives the evolution of suicidal reproduction in mammals. Proceedings of the National Academy of Sciences, 110(44), 17910–17914. 10.1073/pnas.1310691110

Fisher, D. O., Double, M. C., Blomberg, S. P., Jennions, M. D., & Cockburn, A. (2006). Post-mating sexual selection increases lifetime fitness of polyandrous females in the wild. Nature, 444(7115), 89–92. 10.1038/nature05206

Fisher, D. O., Double, M. C., & Moore, B. D. (2006). Number of mates and timing of mating affect offspring growth in the small marsupial *Antechinus agilis*. Animal Behaviour, 71(2), 289–297. 10.1016/j.anbehav.2005.03.041

Fisher, D. O., & Salguero-Gómez, R. (2026). Evidence for adaptive explanations of semelparity in animals. Biological Reviews, 1–19. 10.1002/brv.70197

Fitzpatrick, J. L. (2020). Sperm competition and fertilization mode in fishes. Philosophical Transactions of the Royal Society B: Biological Sciences, 375(1813), 20200074. 10.1098/rstb.2020.0074

Foerster, K., Coulson, T., Sheldon, B. C., Pemberton, J. M., Clutton-Brock, T. H., & Kruuk, L. E. B. (2007). Sexually antagonistic genetic variation for fitness in red deer. Nature, 447(7148), 1107–1110. 10.1038/nature05912

Forster, L. (1992). The Stereotyped Behavior of Sexual Cannibalism in Latrodectus-Hasselti Thorell (Araneae, Theridiidae), the Australian Redback Spider. Australian Journal of Zoology, 40(1), 1–11. 10.1071/ZO9920001

Gadgil, M., & Bossert, W. H. (1970). Life Historical Consequences of Natural Selection. The American Naturalist, 104(935), 1–24.

Geritz, S. A. H., Kisdi, É., Meszéna, G., & Metz, J. A. J. (1998). Evolutionarily singular strategies and the adaptive growth and branching of the evolutionary tree. Evolutionary Ecology, 12(1), 35–57. 10.1023/A:1006554906681

Guillet, S., & Vancassel, M. (2001). Dermapteran life-history evolution and phylogeny with special reference to the Forficulidae family. Evolutionary Ecology Research, 3, 441–447.

Gustafsson, L., & Sutherland, W. J. (1988). The costs of reproduction in the collared flycatcher Ficedula albicollis. Nature, 335(6193), 813–815. 10.1038/335813a0

Healy, K., Ezard, T. H. G., Jones, O. R., Salguero-Gómez, R., & Buckley, Y. M. (2019). Animal life history is shaped by the pace of life and the distribution of age-specific mortality and reproduction. Nature Ecology & Evolution, 3(8), 1217–1224. 10.1038/s41559-019-0938-7

Holleley, C. E., Dickman, C. R., Crowther, M. S., & Oldroyd, B. P. (2006). Size breeds success: Multiple paternity, multivariate selection and male semelparity in a small marsupial, Antechinus stuartii. Molecular Ecology, 15(11), 3439–3448. 10.1111/j.1365-294X.2006.03001.x

Hughes, P. W. (2017). Between semelparity and iteroparity: Empirical evidence for a continuum of modes of parity. Ecology and Evolution, 7(20), 8232–8261. 10.1002/ece3.3341

Hurd, L. E. (1989). The importance of late season flowers to the fitness of an insect predator, Tenodera sinensis Saussure (Orthoptera: Mantidae), in an old field community. The Entomologist, 108, 223–228.

Jellyman, D. J. (2022). An enigma: How can freshwater eels (Anguilla spp.) be such a successful genus yet be universally threatened? Reviews in Fish Biology and Fisheries, 32(2), 701–718. 10.1007/s11160-021-09658-8

Karman, L., Genevcius, B. C., Bezerra, W. C., Medrano, M., & Machado, G. (2026). Do contest-related traits evolve faster? A test of the exaggerated evolution hypothesis in a clade of neotropical arachnids. Biology Letters, 22(1), 20250600. 10.1098/rsbl.2025.0600

Karsten, K. B., Andriamandimbiarisoa, L. N., Fox, S. F., & Raxworthy, C. J. (2008). A unique life history among tetrapods: An annual chameleon living mostly as an egg. Proceedings of the National Academy of Sciences, 105(26), 8980–8984. 10.1073/pnas.0802468105

Kindsvater, H. K., Braun, D. C., Otto, S. P., & Reynolds, J. D. (2016). Costs of reproduction can explain the correlated evolution of semelparity and egg size: Theory and a test with salmon. Ecology Letters, 19(6), 687–696. 10.1111/ele.12607

Knoflach, B., & Harten, A. van. (2001). Tidarren argo sp. nov. (Araneae: Theridiidae) and its exceptional copulatory behaviour: emasculation, male palpal organ as a mating plug and sexual cannibalism. Journal of Zoology, 254(4), 449–459. 10.1017/S0952836901000954

Kokko, H. (2021). The stagnation paradox: The ever-improving but (more or less) stationary population fitness. Proceedings of the Royal Society B: Biological Sciences, 288(1963), 20212145. 10.1098/rspb.2021.2145

Kokko, H., & Brooks, R. (2003). Sexy to die for? Sexual selection and the risk of extinction. Annales Zoologici Fennici, 40, 207–219.

Kuntner, M., Kralj-Fišer, S., Schneider, J. M., & Li, D. (2009). Mate plugging via genital mutilation in nephilid spiders: An evolutionary hypothesis. Journal of Zoology, 277(4), 257–266. 10.1111/j.1469-7998.2008.00533.x

Lane, J. E., Boutin, S., Speakman, J. R., & Humphries, M. M. (2010). Energetic costs of male reproduction in a scramble competition mating system. Journal of Animal Ecology, 79(1), 27–34. 10.1111/j.1365-2656.2009.01592.x

Lazenby-Cohen, K. A., & Cockburn, A. (1988). Lek Promiscuity in a Semelparous Mammal, Antechinus stuartii (Marsupialia: Dasyuridae)? Behavioral Ecology and Sociobiology, 22(3), 195–202.

Miller, J. A. (2007). Repeated Evolution of Male Sacrifice Behavior in Spiders Correlated with Genital Mutilation. Evolution, 61(6), 1301–1315. 10.1111/j.1558-5646.2007.00115.x

Mills, S. C., Koskela, E., & Mappes, T. (2012). Intralocus sexual conflict for fitness: Sexually antagonistic alleles for testosterone. Proceedings of the Royal Society B: Biological Sciences, 279(1735), 1889–1895. 10.1098/rspb.2011.2340

Møller, A. P., & Briskie, J. V. (1995). Extra-Pair Paternity, Sperm Competition and the Evolution of Testis Size in Birds. Behavioral Ecology and Sociobiology, 36(5), 357–365.

Monnin, T., & Peeters, C. (1998). Monogyny and regulation of worker mating in the queenless ant *Dinoponera quadriceps*. Animal Behaviour, 55(2), 299–306. 10.1006/anbe.1997.0601

Murphy, G. I. (1968). Pattern in Life History and the Environment. The American Naturalist, 102(927), 391–403. 10.1086/282553

Oli, M. K., & Dobson, F. S. (2003). The relative importance of life-history variables to population growth rate in mammals: Cole’s prediction revisited. The American Naturalist, 161(3), 422–440. 10.1086/367591

Orzack, S. H., & Tuljapurkar, S. (1989). Population Dynamics in Variable Environments. VII. The Demography and Evolution of Iteroparity. The American Naturalist, 133(6), 901–923.

R Core Team. (2025). R: A Language and Environment for Statistical Computing. R Foundation for Statistical Computing. https://www.R-project.org/

Ramos, M., Irschick, D. J., & Christenson, T. E. (2004). Overcoming an evolutionary conflict: Removal of a reproductive organ greatly increases locomotor performance. Proceedings of the National Academy of Sciences, 101(14), 4883–4887. 10.1073/pnas.0400324101

Rankin, D. J., Dieckmann, U., & Kokko, H. (2011). Sexual Conflict and the Tragedy of the Commons. The American Naturalist, 177(6), 780–791. 10.1086/659947

Reznick, D., Travis, J., & Bassar, R. (2026). The ‘big houses-big cars dilemma’: How do we reconcile the prevalence of positive correlations among life history traits? Functional Ecology. 10.1111/1365-2435.70458

Salguero-Gómez, R., Jones, O. R., Jongejans, E., Blomberg, S. P., Hodgson, D. J., Mbeau-Ache, C., Zuidema, P. A., de Kroon, H., & Buckley, Y. M. (2016). Fast–slow continuum and reproductive strategies structure plant life-history variation worldwide. Proceedings of the National Academy of Sciences, 113(1), 230–235. 10.1073/pnas.1506215112

Santos, E. S. A., & Nakagawa, S. (2012). The costs of parental care: A meta-analysis of the trade-off between parental effort and survival in birds. Journal of Evolutionary Biology, 25(9), 1911–1917. 10.1111/j.1420-9101.2012.02569.x

Sasson, D. A., & Ryan, J. F. (2017). A reconstruction of sexual modes throughout animal evolution. BMC Evolutionary Biology, 17(1), 242. 10.1186/s12862-017-1071-3

Schaffer, W. M. (1974a). Optimal Reproductive Effort in Fluctuating Environments. The American Naturalist, 108(964), 783–790.

Schaffer, W. M. (1974b). Selection for Optimal Life Histories: The Effects of Age Structure. Ecology, 55(2), 291–303. 10.2307/1935217

Schwartz, S. K., Wagner, W. E., & Hebets, E. A. (2013). Spontaneous male death and monogyny in the dark fishing spider. Biology Letters, 9(4), 20130113. 10.1098/rsbl.2013.0113

Schwartz, S. K., Wagner, W. E., & Hebets, E. A. (2014). Obligate male death and sexual cannibalism in dark fishing spiders. Animal Behaviour, 93, 151–156. 10.1016/j.anbehav.2014.04.037

Smith, F., & Charnov, E. (2001). Fitness trade-offs select for semelparous reproduction in an extreme environment. Evolutionary Ecology Research, 3, 595–602.

Soulsbury, C. D. (2010). Genetic Patterns of Paternity and Testes Size in Mammals. PLOS ONE, 5(3), e9581. 10.1371/journal.pone.0009581

Tsukamoto, K., Chow, S., Otake, T., Kurogi, H., Mochioka, N., Miller, M. J., Aoyama, J., Kimura, S., Watanabe, S., Yoshinaga, T., Shinoda, A., Kuroki, M., Oya, M., Watanabe, T., Hata, K., Ijiri, S., Kazeto, Y., Nomura, K., & Tanaka, H. (2011). Oceanic spawning ecology of freshwater eels in the western North Pacific. Nature Communications, 2(1), 179. 10.1038/ncomms1174

van Noordwijk, A. J., & de Jong, G. (1986). Acquisition and Allocation of Resources: Their Influence on Variation in Life History Tactics. The American Naturalist, 128(1), 137–142.

Wang, Z. Y., & Ragsdale, C. W. (2018). Multiple optic gland signaling pathways implicated in octopus maternal behaviors and death. The Journal of Experimental Biology, 221(19), jeb185751. 10.1242/jeb.185751

Young, T. P. (1981). A General Model of Comparative Fecundity for Semelparous and Iteroparous Life Histories. The American Naturalist, 118(1), 27–36.

Young, T. P. (1990). Evolution of semelparity in Mount Kenya lobelias. Evolutionary Ecology, 4(2), 157–171. 10.1007/BF02270913

